# Transbilayer Coupling of Lipids in Cells Investigated by Imaging Fluorescence Correlation Spectroscopy

**DOI:** 10.1101/2022.01.06.475300

**Authors:** Nirmalya Bag, Erwin London, David A. Holowka, Barbara A. Baird

**Affiliations:** Department of Chemistry and Chemical Biology, Cornell University, Ithaca, NY; Department of Biochemistry and Cell Biology, Stony Brook University, Stony Brook, NY

## Abstract

Plasma membrane hosts numerous receptors, sensors, and ion channels involved in cellular signaling. Phase separation of the plasma membrane is emerging as a key biophysical regulator of signaling reactions in multiple physiological and pathological contexts. There is much evidence that plasma membrane composition supports the co-existence liquid-ordered (Lo) and liquid-disordered (Ld) phases or domains at physiological conditions. However, this phase/domain separation is nanoscopic and transient in live cells. It is recently proposed that transbilayer coupling between the inner and outer leaflets of the plasma membrane is driven by their asymmetric lipid distribution and by dynamic cytoskeleton-lipid composites that contribute to the formation and transience of Lo/Ld phase separation in live cells. In this Perspective, we highlight new approaches to investigate how transbilayer coupling may influence phase separation. For quantitative evaluation of the impact of these interactions, we introduce an experimental strategy centered around Imaging Fluorescence Correlation Spectroscopy (ImFCS), which measures membrane diffusion with very high precision. To demonstrate this strategy we choose two well-established model systems for transbilayer interactions: crosslinking by multivalent antigen of immunoglobulin E bound to receptor FcεRI, and crosslinking by cholera toxin B of GM1 gangliosides. We discuss emerging methods to systematically perturb membrane lipid composition, particularly exchange of outer leaflet lipids with exogenous lipids using methyl alpha cyclodextrin. These selective perturbations may be quantitatively evaluated with ImFCS and other high-resolution biophysical tools to discover novel principles of lipid-mediated phase separation in live cells in the context of their pathophysiological relevance.

## The plasma membrane maintains lipid asymmetry across outer and inner leaflets

The plasma membrane of eukaryotic cells is asymmetric in lipid composition and resulting biophysical and functional properties ^1–3^. As demonstrated for erythrocytes and consistent with results for nucleated cells ^4^, the profile of lipid head group and acyl chain composition differs across leaflets. While glycosphingolipids (GSL) and sphingomyelins (SM) are primarily present in the outer leaflet, phosphatidylethanolamines (PE), phosphatidylserines (PS), and phosphatidylinositol phosphates (PIPs) localize to the inner leaflet. Phosphatidylcholines (PC) are present in both leaflets, and cholesterol flips rapidly between leaflets ^5^. The outer leaflet lipids generally have more saturated chains than the inner leaflet yielding higher chain order and effective viscosity in the former, as shown by measurements of membrane order and diffusion ^4, 6, 7^. The inner leaflet phospholipids have a net negative charge while the outer leaflet phospholipids are zwitterionic. This asymmetry is maintained by well-regulated systems of enzymes and transporters, including flippases, floppases, and scramblases ^3^. In addition to receptors and other membrane proteins that participate in transmembrane function, the extracellular glycocalyx layer encases the outer leaflet while intracellular cortical cytoskeleton apposes and connects to the inner leaflet ^8^. The biophysical and biochemical asymmetry of these two leaflets along with the membrane proximal non-lipid layers are crucial for cellular activities ^2^.

## Phase-like properties of plasma membanes

Lipid-based phase behavior within the plasma membrane is one of the biophysical properties most heavily studied over the last decades ^9–11^. Early indications from biochemical analyses of detergent-resistant membrane fractions ^12^ and recent high-resolution spectroscopic ^13–17^ and super-resolution microscopic ^18–21^ measurements are consistent with the presence of nanoscopic phase-like organization in the steady-state plasma membrane of resting and stimulated cells. Although remaining a subject of controversy, the origin and maintenance of such organization is likely to be driven by multiple, concurrent mechanisms ^11, 22, 23^. We broadly define this phase-like organization as co-existing liquid-ordered like (Lo-like) and liquid-disordered like (Ld-like) regions. The physical properties of Lo- and Ld-like nanodomains resemble those of Lo and Ld phases in artificial membranes of defined lipid composition ^24^, which have also been used to describe microscopic phase separation in cell-derived giant plasma membrane vesicles (GPMVs) ^25, 26^. In cells, the Lo-like nanodomains, which are individually heterogeneous, are roughly grouped together and colloquially referred to as “lipid rafts”. Interestingly, symmetric lipid bilayers mimicking outer-leaflet lipid composition (SM (or saturated PC)/unsaturated PC/cholesterol) phase-separate readily, but the same is not true for symmetric bilayers mimicking inner leaflet composition (PE/PS/PIPs/cholesterol) ^27^. However, inner-leaflet lipid probes are observed to partition preferentially into phase-separated GPMVs ^28^. Thus, transbilayer coupling in plasma membranes appears to be a mechanism influencing phase-like organization in both leaflets ^15, 29, 30^.

## Transbilayer interactions modulate phase-like properties in plasma membrane leaflets

Transmembrane (TM) proteins are obvious candidates for mediating transbilayer interactions in cells. TM proteins can coordinately modulate both leaflets through multiple possible processes including line tension, hydrophobic mismatch, and coalescence of pre-existing nanodomains as the result of protein clustering ^31^. A well-established example for the last of these is transbilayer reorganization caused by antigen (Ag) crosslinking of the high affinity receptor, FcεRI, for immunoglobulin E (IgE), which initiates cellular signaling at the earliest stage of allergic responses. The nanoclusters of Ag-crosslinked IgE-FcεRI locally stabilize Lo-like nanodomains,^32^ which may register across leaflets (Fig. 1, Interaction type 1). This model is extensively tested by theory ^33^, simulation ^34^, and experiments ^18, 35–37^ in mast cell types and also extends to other immunoreceptor signaling systems ^19, 38–40^.

**Figure 1.**
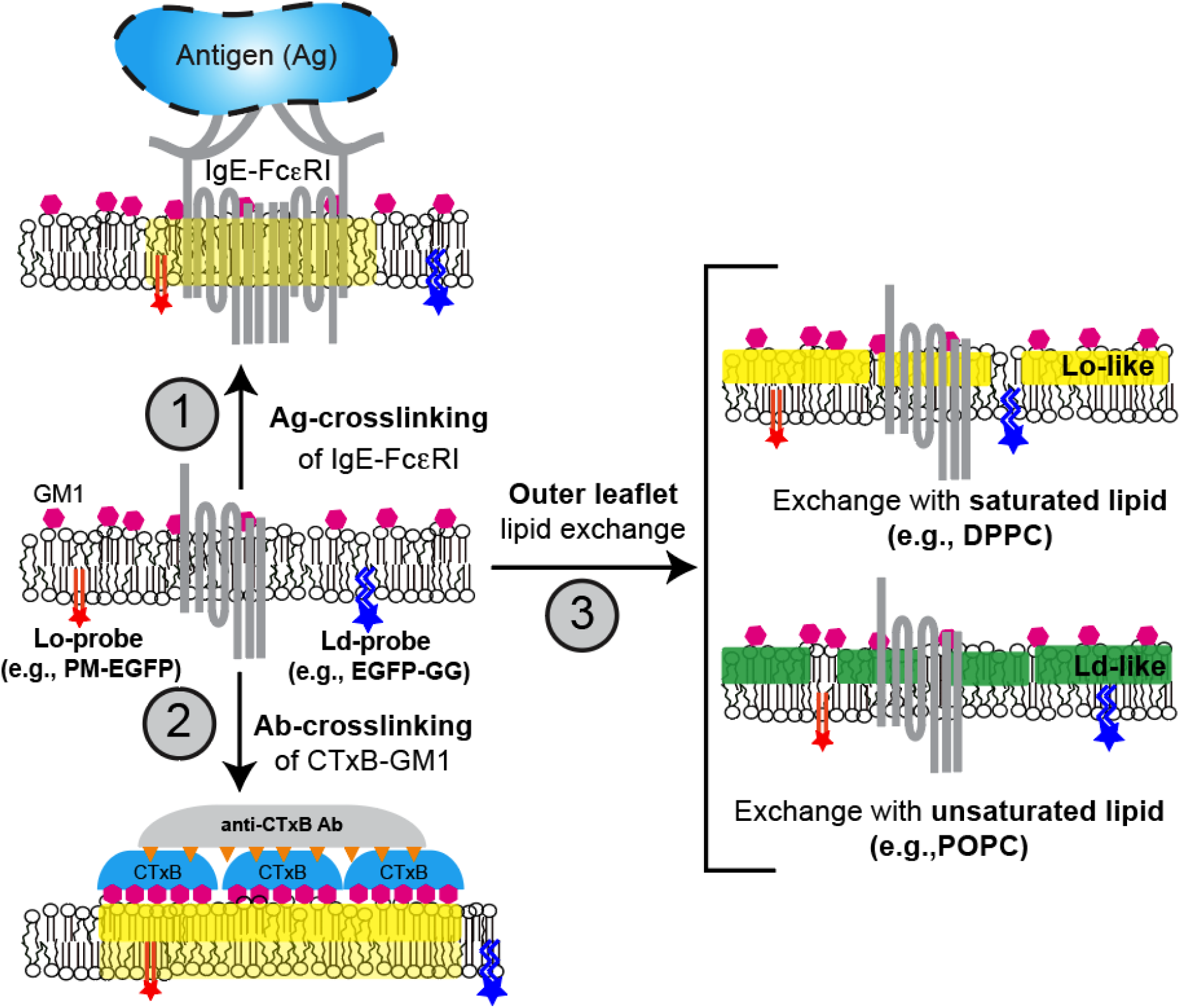
Modalities of ‘outside-in’ transbilayer coupling. 1) Cross-linking of immunoglobulin E (IgE) sensitized transmembrane (TM) receptor FcεRI by extracellular antigen (Ag) may stabilize the Lo-like phase (yellow shade) in both outer and inner leaflets because the receptor spans both leaflets. 2) Cross-linking of pentameric cholera toxin B (CTxB)/GM1 ganglioside complex by anti-CTxB antibodies may stabilize Lo-like phase separation in the outer leaflet which then induces Lo stabilization in the inner leaflet. 3) Outer leaflet lipid exchange (LEX) with exogenous Lo-promoting, saturated lipid (yellow, top) or Ld-promoting, unsaturated lipid (green, bottom). These outer leaflet alternations of phase-specific lipid composition may drive corresponding phase stabilization or destabilization in the inner leaflet.

Based upon simulations and model membrane studies, lipid-mediated transbilayer coupling mechanisms have also been proposed^41, 42^. The definition of lipid-based transbilayer coupling varies in the literature depending on the context. Here we define it as a process by which changes in the lipid phase organization of one leaflet induce changes in the other leaflet ^43^. Both ‘outside-in’ and ‘inside-out’ mechanisms are proposed. Prominently suggested for an ‘outside-in’ mechanism is interdigitation of outer leaflet, long-chain sphingolipids ^44, 45^ or ceramides ^46, 47^ into the inner leaflet, such that ordering in the outer layer is conferred upon the inner layer. As has been evaluated in live cells, clustering of outer-leaflet, Lo-preferring components appear to induce Lo-like domains in the inner leaflet (Fig. 1, Interaction mode 2). Specifically, immobilization of GM1, an outer-leaflet, Lo-preferring glycosphingolipid, crosslinked extensively by cholera toxin B (CTxB) was shown to cause detectable stabilization of Lo-like nanodomains in the inner leaflet ^48^. Lipid interdigitation may also be actively manipulated by actin in an ‘inside-out’ mode ^15, 49^. For example, dynamic actin monomers appear to attach to and stabilize nanoclusters of interdigitating, long-chain PS lipids in the inner leaflet, which then interact with long-chain outer leaflet lipids and lipid-anchored proteins (e.g., glycosylphosphatidylinositol (GPI)-anchored proteins) to create registered Lo-like nanodomains in both leaflets ^15^. Computer simulations on this system indicated that such transbilayer coupling by immobilization of one of the long-chain components (PS or GPI) occurs by recruitment of Lo-preferring components without requiring pre-existing lipid-based phase separation in either leaflet ^15^.

Although some of the above studies on transbilayer coupling were carried out on live cells, these measurements typically require specialized microscopes and difficult sample preparation, which limit their wide applicability. This is a reason why many models of transbilayer coupling are tested by simulation and experiments on asymmetric model membranes. For example, Chiantia and London measured diffusion (by confocal fluorescence correlation spectroscopy (FCS)) of lipid probes and membrane order in both leaflets of asymmetric lipid bilayers to show that lipid-based transbilayer coupling depends on both saturation and acyl chain length of the lipids ^44^. More recently, the London group developed a straightforward strategy to alter the outer leaflet lipid composition in live cells by exchange with selected exogenous lipids,^50, 51^ opening another window to test how outer-leaflet lipid composition influences the organization of the inner leaflet (Fig. 1, Interaction mode 3). They showed that membrane order of GPMVs isolated after lipid exchange in cells (most of the outer leaflet lipids are replaced) decreases or increases, compared to unexchanged conditions, with unsaturated lipids (Ld-promoting) or saturated lipids (Lo-promoting), respectively ^52^. However, the specific impact of this outside-in mechanism, i.e, extent of effect caused by outer leaflet lipid exchange on inner leaflet properties, was not quantified in these studies.

As described above, both outside-in and inside-out approaches have been developed to create or modulate transbilayer coupling in model membranes and live cells. However, still lacking are readily accessible experimental platforms that precisely quantify the degree of transbilayer interactions occurring under a range of conditions. Ease of application and rigorous quantification are necessary to evaluate the importance of these intra-membrane interactions in cellular functions.

## Diffusion of phase-selective lipid probes reflects underlying phase-like properties of plasma membrane leaflets

Work in our and other laboratories has shown that diffusion of membrane components can be measured to monitor interactions within each leaflet as well as coupling between the leaflets. This has been particularly explored in the context of phase-dependent behaviors ^4, 7, 44, 53^. Microscopic phase separation in synthetic giant unilamellar vesicles and GPMVs is readily observed by diffraction-limited fluorescence imaging ^28, 54^. More recently, nanoscopic domains in these model membranes were directly imaged by cryo-electron microscopy ^55^ and cryo-tomography ^56^. However, researchers have been generally unsuccessful in directly imaging these nanodomains in live cells with conventional microscopy. Some groups characterized their presence at nanometer scale with fluorescence resonance energy transfer (FRET) using probes independently shown to partition into Lo or Lo domains in phase-separated model membranes.^57, 58^ Also, super-resolution microscopy has systematically correlated localization of Lo- or Ld-preferring probes within their corresponding regions as identified by other markers ^18, 19, 59^.

Fluorescence recovery after photobleaching (FRAP), FCS, and single particle tracking (SPT) have been developed to measure diffusion in live cells.^60^ Among them, FRAP ^61^ and FCS ^62^ have had limited success detecting underlying phase-like organization using diffraction-limited microscopy. Efforts to improve detection led to the development of spectral ^16, 63^ and spatial analysis modalities, including, spot variation modalities of FCS ^64^ and FRAP ^65^, image mean-squared-displacement ^66^, and pair correlation functions ^67, 68^, which are typically applied on diffraction-limited measurements and are extrapolated to gain nanoscopic information indirectly. The Eggeling group directly evaluated phase-like organization in resting live cells using stimulated emission depletion (STED)-FCS on Lo-preferring GPI-anchored protein probes and a range of synthetic Lo and Ld-preferring lipid probes labelled with organic fluorophores in outer membrane leaflet ^13^. They found diffusion behavior of these probes to be consistent with the presence of lipid-dependent nanoscopic entities in which Lo-preferring lipid probes, but not their Ld-preferring counterparts, are confined for several milliseconds. These nanoscopic entities appear to be Lo-like nanodomains because they specifically confine Lo-preferring probes. Other groups developed alternative modalities of nanoscopic FCS measurements and showed similar confinement of Lo-preferring probes, although the average confinement times and length scales of confinement derived from analysis of these measurements vary widely ^69, 70^. In parallel, several groups, notably Kusumi and colleagues, developed highly sophisticated microscopes and specialized fluorescent probes for SPT, incorporating robust analysis to evaluate diffusion of single lipids ^71, 72^. These various spectroscopic analyses provide substantial evidence for existence of transient, nanoscopic Lo-like regions which preferentially confine Lo-preferring, saturated lipid probes. However, physical characterization of the nanodomains (e.g., size, lifetime, mobility) extracted from the measured diffusion properties of these probes is challenging and varies broadly.

Although detailed information has come from these high-resolution tools, they are typically technically demanding which significantly limits their application in wide ranging biophysical problems of plasma membranes. Readily accessible, complementary methods are called for. As described below, camera-based imaging FCS is proving to advance this purpose, requiring only recording of time-lapse (diffraction-limited) fluorescence movies of a sample plane by a fast camera ^73^. The computation of diffusion coefficients and several other spatial fluctuation parameters from this time-lapse movie is done by software, which has been developed, is actively maintained, and is freely available from the Wohland group^74^.

## Subtle changes in membrane diffusion are quantified by statistically-robust Imaging FCS

Imaging FCS is realized in both total internal reflection fluorescence (TIRF) and light sheet microscopes equipped with an EMCCD or sCMOS camera ^75^. Here, we showcase the results from TIRF-based imaging FCS (ImFCS), also known as imaging TIR FCS (ITIR-FCS) ^76^, which is most readily applied to examine the ventral plasma membrane. We show how diffusion measurements for multiple probes with different preferences for lipid phase and protein interactions reveal cell membrane heterogeneity (Fig. 2). Light sheet-based ImFCS can be employed to illuminate plasma membrane properties in live organisms ^77^. Below we describe the ImFCS set up in our laboratory; we refer to ^78^ for a comprehensive review of this approach.

**Figure 2.**
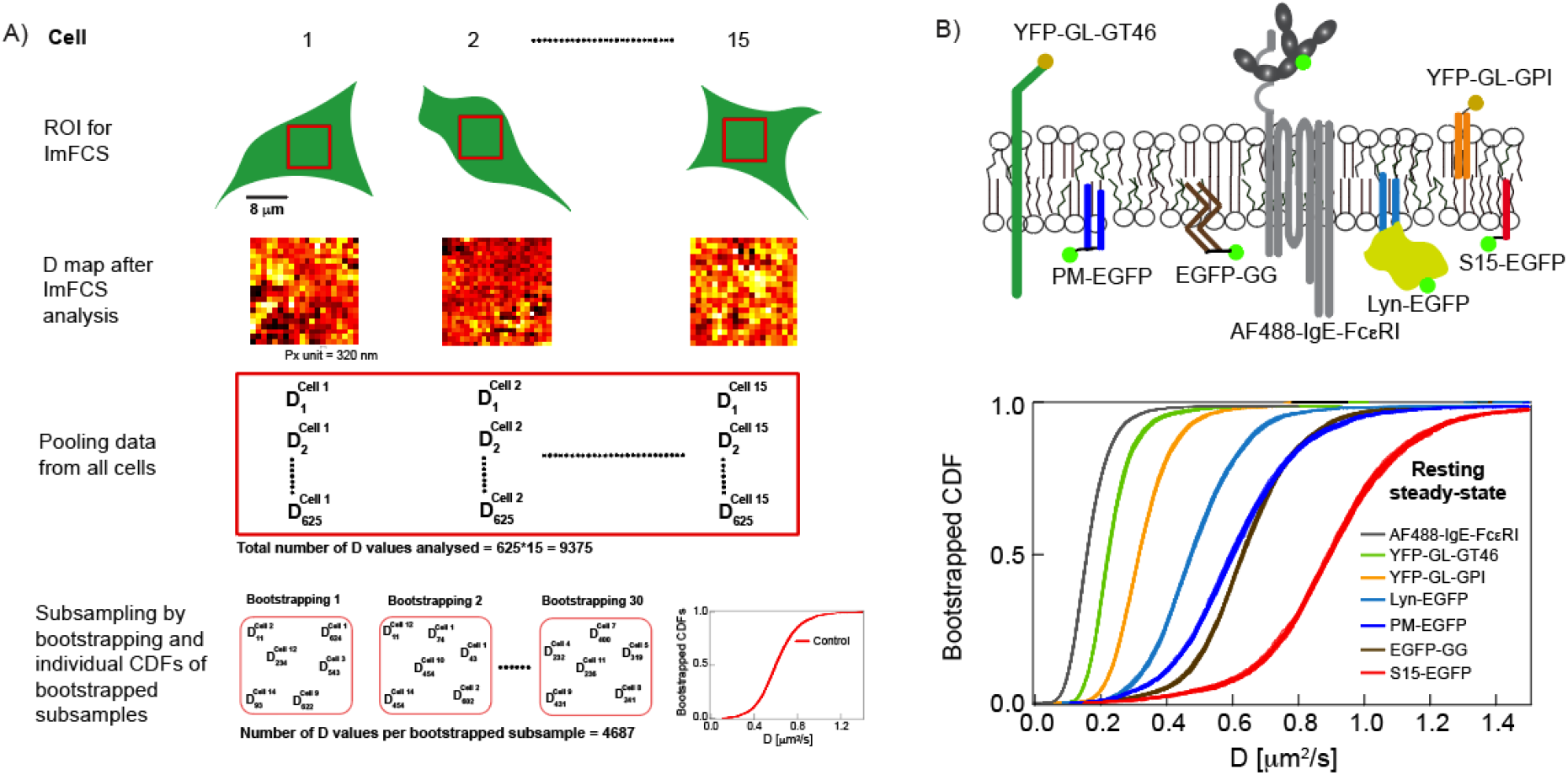
Demonstration of Imaging FCS data analysis (A) and wide range of diffusion coefficients shown by membrane probes depending on their localization and membrane attachment (B). A) ImFCS analysis of time-lapse TIRF movies of a chosen regions of interest (ROI) yields spatial maps of diffusion coefficient (*D*). Here we show a 25×25 Px unit ROI (Px unit = 320×320 nm^2^) and representative *D* map of same dimension for EGFP-GG expressed in RBL cells. This ImFCS analysis yields 625 *D* values from one cell. To improve data statistics and precison, the *D* values from multiple cells are pooled (illustrated as red rectangle). The pooled data is subsampled by bootstrapping and the bootstrapped *D* sub-samples are translated into cumulative distribution functions which are overlaid to check for heterogeneity across all data. B) The transmembrane probes, yellow fluorescent protein (YFP)-tagged GT46 (YFP-GL-GT46) ^117^ and alexa fluor 488 (AF488) tagged IgE-FcεRI complex (AF488-IgE-FcεRI) ^118^, are Ld-preferring. The outer leaflet lipid probe, YFP-taqged Glycosylphosphatidylinositol (YFP-GL-GPI)^118^, is Lo-preferring. Among the inner leaflet probes, enhanced green fluorescent protein (EGFP)-tagged palmitoyl and myristoyl anchor (PM-EGFP) ^118, 119^ and Lyn kinase (Lyn-EGFP) ^118^ are Lo-preferring while EGFP-tagged gernyl-gernyl anchor (EGFP-GG)^118, 119^ and myristoyl anchor (S15-EGFP)^120^ are Ld-preferring.

ImFCS measures diffusion of probes in both leaflets of the ventral plasma membrane with very high precision (Fig. 2A) ^7, 37^. The probes can be tagged with common fluorophores such as genetically-encoded fluorescent proteins or organic fluorophores. The combination of z-sectioning by TIRF illumination (100 nm) and xy-sectioning as defined by an array of pixel units (Px unit = 320×320 nm^2^ in our laboratory set up) on the camera chip provides a sufficiently small observation area to allow autocorrelation function (ACF) analysis of the temporal fluorescence fluctuations for each Px unit. The diffusion coefficient (*D*) extracted from this pixel-specific analysis averages over the acquisition time (typically 80,000 frames at 3.5 msec/frames or 280 seconds in our laboratory set up). One time-lapse movie contains several hundreds Px units across the imaged membrane area (roughly 8×8 μm^2^), and thereby several hundreds parallel diffusion measurements for one cell. These provide a spatial map and an ensemble of *D* values that can be plotted as a distribution or simply averaged for that cell. Increasing the ensemble size by combining data from multiple cells and then subsampling multiple times by bootstrapping to minimize the effect of unusual outliers, underlies the high confidence level of *D* values determined that are specific to each probe ^37^.

We recently employed ImFCS to measure diffusion of various probes in rat basophilic leukemia (RBL) cells, an established mast cell model (Fig. 2B) ^7^. Compiling ~10,000 *D* values from about 15 cells for each probe in the form of a cumulative distribution function (CDF) allows distinctive curve shapes and averaged values of *D* (*D_av_*) to be determined. Our bootstrapping approach allows us to evaluate the precision of the CDF and *D_av_* of a probe under a given condition, as represented by the thickness of the CDF curve. This analysis showed the error in our *D_av_* values to be <1%. We demonstrated the sensitivity of ImFCS by comparing diffusion properties of inner leaflet, outer leaflet, and TM probes in the plasma membrane of resting RBL cells (Fig. 2B) ^7^. The diffusion of these structurally distinct membrane probes in the plasma membrane varies significantly depending on the membrane attachment of the probes and their potential protein interactions. The CDF of *D* values (after bootstrapping-based sub-sampling) characterizes a given probe in terms of position (*D_av_*) and shape (heterogeneity of *D* values), which is well fitted by a one- or two-component Gaussian model ^7^. In general, lipid-anchored probes diffuse faster than TM probes (e.g., *D_av_* (YFP-GL-GPI) > *D_av_* (YFP-GL-GT46)), and outer membrane lipid probes diffuse slower than their inner leaflet counterparts (e.g., Lo-preferring: *D_av_*(YFP-GL-GPI) < *D_av_*(PM-EGFP)) (Fig. 2B). Among the lipid probes in the inner leaflet, Lo-preferring probes diffuse slower than Ld-preferring probes (e.g., *D_av_*(PM-EGFP) < *D_av_*(EGFP-GG)). These results are consistent with previously measured differences in the lateral diffusion and order parameters of Lo vs Ld regions in inner and outer leaflets ^4, 6, 61^. However, although it was previously possible to measure large differences in *D* values among membrane probes, small differences (e.g., PM-EGFP, *D_av_*=0.62 ± 0.002 μm^2^/sec vs EGFP-GG, *D_av_*=0.64 ± 0.002 μm^2^/sec; Fig. 2B) were not previously discernable ^61, 79^. The exceptional level of sensitivity, based on the data statistics of diffusion measurements by ImFCS, allows small differences in the plasma membrane organization, as experienced by diffusing probes, to be elucidated. This sensitivity opened the door to delineating the subtle modulation of Lo/Ld-like organization after Ag-crosslinking of IgE-FcεRI, which initiates signal transduction in these cells ^37^, as described in the following section.

It is broadly recognized that inter-leaflet asymmetry within the plasma membrane and mechanisms for transbilayer coupling are essential for initiating transmembrane signaling as well as for maintaining cell homeostasis. Because coupling is manifest as changes in one leaflet caused by perturbation of the other leaflet, this can result in changes in the diffusion properties of membrane components in both leaflets. Such induced diffusional changes may be subtle, but precise measurements with ImFCS enable small differences to be discerned. Together with defined perturbations, ImFCS holds great promise for delineating principles of transbilayer coupling in live cells. In following sections, we focus on the effects of outside-in coupling modes on the Lo/Ld phase-like organization of the inner leaflet by measuring diffusional changes of probes PM-EGFP (Lo-preferring) and EGFP-GG (Ld-preferring) (Figs. 1 and 2B).

## Antigen (Ag)-crosslinking of transmembrane IgE-FcεRI locally stabilizes Lo/Ld-like phase separation in the membrane inner leaflet

As exemplified by immune receptors, clustering of TM proteins by extracellular multivalent ligands can modulate Lo/Ld-like organization across the plasma membrane. Crosslinking of IgE-FcεRI by multivalent Ag to stimulate RBL cells results in largely immobilized clusters of these TM receptors ^37, 80^. ImFCS measurements before and after crosslinking IgE-FcεRI showed that PM-EGFP diffusion slows (*D* CDF shifts left), whereas EGFP-GG diffusion becomes faster (*D* CDF shifts right) (Fig. 3A)^37^. These results extend previous super-resolution imaging and confirm that the lipid environment in the inner leaflet is changed around the clustered IgE-FcεRI: Lo/Ld phase-like separation is stabilized. Super-resolution imaging showed nanoscale co-localization of the analogous fluorescent palmitate/myristoylate (PM) probe with IgE-FcεRI only after Ag-crosslinking, whereas the analogous fluorescent geranylgeranyl (GG) probe does not co-localize^18^. The slower diffusion of PM-EGFP measured with ImFCS confirms nanoscopic stabilization of an Lo-like phase surrounding the clustered receptors and further indicates a distinctive Ld-like phase distally^37^. The full set of diffusion measurements in this study delineated the lipid-based and protein-based interactions that synergize during Ag-stimulated transmembrane signaling. Together they provide compelling quantitative evidence that Ag-stimulated transmembrane signaling is mediated by lipid-based coupling of Lyn kinase (lipid-anchored to the inner leaflet) with clustered IgE-FcεRI with simultaneous exclusion of a TM phosphatase ^37^. Our ImFCS measurements also showed that Ag-crosslinking slows the diffusion of an outer leaflet Lo-preferring probe, YFP-GL-GPI ^37^, suggesting similar Lo/Ld stabilization also occurs around clustered IgE-FcεRI in the outer leaflet. Distinctive phase separation in the outer leaflet remains to be tested with an outer leaflet Ld-preferring probe. Because of the marked differences in lipid composition and ordering in outer vs inner membrane leaflets, it is possible that Lo/Ld rearrangements after antigen-clustering of IgE-FcεRI occurs differentially in these two leaflets.

**Figure 3.**
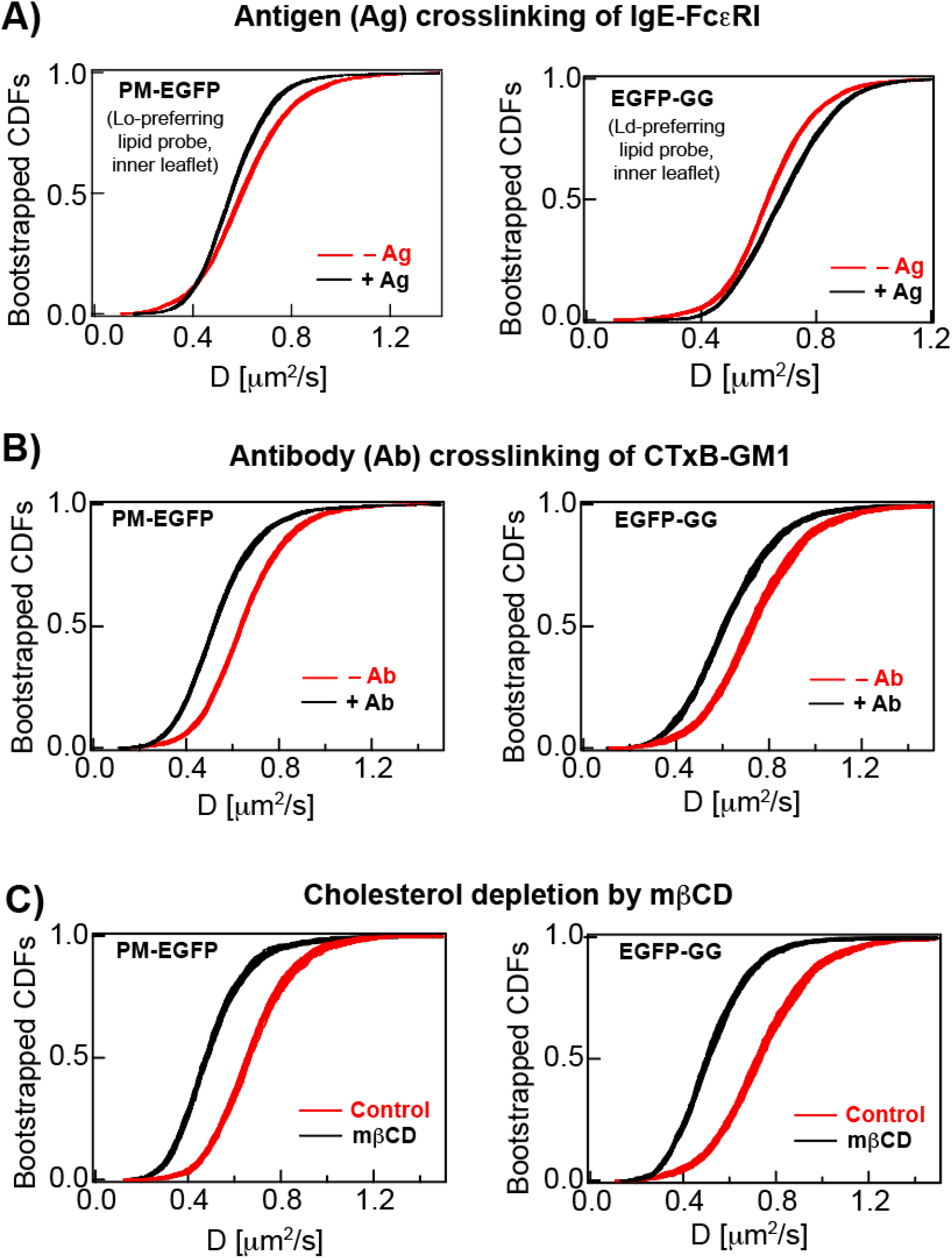
Changes of diffusion for Lo-preferring (PM-EGFP) and Ld-preferring (EGFP-GG) lipid probes, after A) crosslinking of IgE-FcεRI complex by multivalent antigen (Ag), B) crosslinking of cholera toxin B– GM1 (CTxB-GM1) complex by polyclonal antibody (Ab), and C) cholesterol extraction by 5 mM mβCD at 37°C for 30 min.

## Antibody (Ab)-crosslinking of glycosphingolipid complex CTxB-GM1 globally affects inner leaflet diffusion properties

To evaluate lipid-mediated transbilayer coupling, we used ImFCS to measure the impact on diffusion of probes in the inner leaflet caused by crosslinking of GM1, an outer leaflet, Lo-preferring glycosphingolipid ^81^. Cholera toxin B (CTxB) is commonly used as an exogenous ligand that binds to as many as five copies of GM1 (Fig. 1, mode 2). Crosslinking CTxB-GM1 complexes with specific antibodies was shown to coalesce and stabilize Lo-like nanodomains in the membrane outer leaflet in live cells, as reflected by redistribution of Lo- and Ld-preferring probes ^82^. These measurements, made with single-molecule near-field scanning optical microscopy, provided direct evidence for the connectivity of nanoscale Lo-like domains in the outer leaflet, such that these can be coalesced by clustering Lo-preferring constituents. The Kusumi group evaluated Ab-crosslinked CTxB-GM1 clusters with SPT using their ultra-high-speed video camera to test for induced reorganization in the inner leaflet ^48^. They observed transient but significant co-localization of inner leaflet Lo-preferring components, including lipid probes and lipid-anchored protein probes. They further observed similar co-localization on the inner leaflet after Ab-crosslinking GPI-anchored CD59, an outer leaflet, Lo-preferring proteolipid. These observations indicate that coalescence of Lo-preferring components in the outer leaflet can register Lo-preferring components in the inner leaflet, possibly by interdigitation of long, saturated lipids.

We evaluated effects of glycosphingolipid stabilization in our RBL cell experimental system, where diffraction-limited TIRF microscopy showed microscopic clusters of fluorescently labeled CTxB-GM1 after crosslinking by Ab. Then, using nonfluorescent Ab-GM1-CTxB, we evaluated the properties of inner leaflet probes PM-EGFP and EGFP-GG, and found these exhibited no obvious clustering. This observation may be explained by the evident miscibility of the inner leaflet lipid composition ^27^, such that any phase segregation is sub-diffraction before or after clustering an outer leaflet lipid component. Effects on the inner leaflet caused by perturbations in the outer leaflet require more sensitive measurements, which can be achieved with ImFCS.

ImFCS diffusion measurements of inner leaflet probes before and after Ab-crosslinking of CTxB-GM1 showed a 16% decrease in *D_av_* for PM-EGFP (Fig. 3B). This is in the same direction as the effect on PM-EGFP diffusion caused by Ag-crosslinking IgE-FcεRI (8% decrease; Fig. 3A) and consistent with the Kusumi group finding that transbilayer coupling affects phase segregation in the inner leaflet ^48^. The larger diffusional decrease we observe for Ab-CTxB-GM1 compared to Ag-IgE-FcεRI may reflect differences in the mediator of transbilayer coupling (lipids vs TM protein). However, other differences between Ab-CTxB-GM1 and Ag-IgE-FcεRI systems include the degree of clustering under the experimental conditions evaluated; this and other possibilities, such as actin-engagement ^18, 83^ or effects on membrane curvature^84^ remain to be investigated. Interestingly, we also observed similar decrease of EGFP-GG’s diffusion after Ab-crosslinking of CTxB-GM1 (Fig. 3B), which is a direction different from the diffusional increase we observed for this probe with Ag-IgE-FcεRI (Fig. 3A). These contrasting results underscore differences that depend on the means of transbilayer coupling. They further suggest that extensively clustering Lo-preferring lipids in the outer leaflet can cause global changes in the phase-like behavior in both the outer and inner leaflets, such that, for example, the inner leaflet also becomes more ordered overall.

## Reduction in plasma membrane cholesterol globally slows ImFCS-measured diffusion of inner leaflet probes

A common strategy to study membrane phase separation is by monitoring dependence on the abundance of cholesterol, a key component component of Lo domain formation^85, 86^. Cholesterol chelation by methyl-β-cyclodextrin (mβCD) is often used to reduce cholesterol content in plasma membranes and thereby perturb phase-like properties. This method can lead to insights into roles of membrane phases, if the results are carefully interpreted and the uncertainties associated with mβCD treatments are considered ^87^. For example, there is no general consensus on the degree to which cholesterol is extracted from each leaflet, given the high flip-flop rate of cholesterol across the bilayer. Moreover, conventional biochemical assays only provide total cholesterol amounts in cells and not specifically from the plasma membranes^88^. Acute cholesterol reduction in live cell membranes by mβCD is further complicated by any alterations in cholesterol-dependent cellular processes. To avoid many of the issues related to cholesterol extraction, a sterol substitution strategy can be used, in which sterol levels are maintained while the ability of sterol to supported ordered domain formation is varied ^89^.

We used ImFCS to evaluate mβCD-induced changes in membrane heterogeneity and to compare with other mechanisms that alter Lo/Ld-like properties (Fig. 1). We found that diffusion of both inner leaflet probes, PM-EGFP and EGFP-GG, slows (~25% decrease) after mβCD-extraction of cholesterol (Fig. 3C). Because cholesterol interactions are integral to formation of Lo phases, cholesterol reduction might be expected to make plasma membrane less ordered and more fluid ^62, 90^. However, some previous live cell studies ^61, 91, 92^ indicate that cholesterol extraction can yield a more ordered state, which may be due to formation of gel-like nanodomains, or to alteration of membrane trafficking in the cell. Our ImFCS measurements demonstrate that cholesterol extraction impacts the inner leaflet of the plasma membrane globally.

These early results raise interesting questions to pursue in the future. These involve understanding better the reorganization of residual cholesterol after extraction and the interactions of cholesterol with natural and saturated phospholipids (e.g., SM and DPPC) and phosphatidylserine (PS), which are major cholesterol-interacting plasma membrane lipids ^93^. Both of these phenomena probably depend on the other lipids present, which can be evaluated by exchange of endogenous outer leaflet lipids with selected exogenous lipids ^50, 51^ (see following section). For example, introducing a very high concentration of DPPC in the outer leaflet might cause reorganization of cholesterol across leaflets, such as a net increase of cholesterol in the outer leaflet. This transbilayer movement of cholesterol will likely modify interactions among the remaining constituents of both leaflets and lead to a new plasma membrane steady-state. In general, the redistributed interactions of cholesterol in the plasma membrane after a particular perturbation (such as Ag-IgE-FcεRI, Ab-CTxB-GM1, mβCD, or lipid exchange) must be better understood to interpret the local and global changes measured by ImFCS and other experimental techniques. We expect that effective cholesterol biosensors will be valuable for quantifying cholesterol content in each leaflet of the plasma membrane ^94–98^.

## Exchange of endogenous outer leaflet lipids with exogenous lipids is a new approach for examining transbilayer interactions

The outer leaflet lipid composition may be broadly categorized into three components: saturated acyl chains, unsaturated acyl chains, and cholesterol ^99^. These three components are minimally required to form Lo/Ld phase separation in model membranes. The properties of the individual phases and the corresponding phase diagram depends on structural details of the lipids. A useful strategy for examining lipid-based transbilayer coupling would be to change the composition of any of the three lipid types in the outer leaflet and monitor how that impacts in the inner leaflet. As described in the previous section, decreasing or increasing cholesterol content in live cells with mβCD is used commonly to modulate phase-like behavior ^87^ and consequent effects on transmembrane signaling have been observed ^100^.

In addition to other complicating factors related to cholesterol extraction ^87^, mβCD does not specifically extract cholesterol from the outer leaflet due to rapid flip-flop rate of cholesterol between leaflets ^5^. In contrast, flip-flop rate of phospholipids is very slow ^101^, making it possible to modulate phospholipid and sphingolipid composition specifically in the outer leaflet and monitor effects on transbilayer interactions. Furthermore, altering phospholipid and sphingolipid allows investigation of the role of specific structural features of these lipids in domain formation and protein interactions ^43^.

Li et al. recently presented a lipid exchange (LEX) method that specifically replaces native outer-leaflet phospholipids and sphingolipids with selected exogenous lipids (with saturated or unsaturated acyl chains) in both model vesicles and live cells ^50, 51^. Briefly, the exogenous lipid (lipid_ex_) is first complexed with methyl alpha cyclodextrin (mαCD) followed by addition of this lipid_ex_/mαCD complex directly to the synthetic vesicle or live cell sample. This treatment results in an outer leaflet composition having up to ~80-100% of lipid_ex_ as evaluated in terms of sphingomyelin release using mass spectrometry and other methods ^52^. Lipid flip-flop rate is known to be very slow ^101^: in some cases lipid_ex_ was found to remain in the outer leaflet for several hours ^52^. Importantly, mαCD does not change cholesterol content of the plasma membrane ^102^. The value of the LEX method includes its capacity to change the phase-like organization of the outer leaflet specifically. Indeed, the London group used LEX to create synthetic asymmetric lipid vesicles to test the connection between phase separation and transbilayer interactions ^29, 103^. Their work with this model system confirmed that each leaflet can influence the Lo/Ld phase state of the apposing leaflet. With asymmetric vesicles they further identified particular inner leaflet compositions that suppress phase separation in outer leaflets having lipid compositions that undergo phase separation when present in symmetric vesicles, as well as outer leaflet compositions that induce inner leaflet ordered domain formation. These results with model membranes support the possibility of an entirely lipid-based mechanism in cells where asymmetric inner and outer leaflets interact to modulate phase-like properties. The London group also evaluated nanoscopic phase separation in GPMVs derived from RBL cells at physiological temperature (37°) using a FRET assay ^26^. The GPMVs were collected before or after outer leaflet LEX with either Lo-promoting lipid_ex_ brain sphingomyelin (bSM) or Ld-promoting lipid_ex_ POPC. They observed disappearance of Lo-like nanodomains at 37° for GPMVs collected after POPC (but not bSM) exchange. Thus, changes in lipid compositional asymmetry may regulate the Lo/Ld-like properties in live cell membranes. In addition, in a recent study the London lab found that a loss of lipid asymmetry in GPMVs, which may occur at some point during signal transduction in cells, can induce ordered domain formation ^104^

Together, these results support the view that phase-like organization in the plasma membranes depends on the lipid composition present. However, this interpretation is an extrapolation from model membranes, either synthetic or GPMVs that lack a cytoskeleton and may have altered asymmetry compared to parental cells. Nonetheless these model studies clearly demonstrate wide applicability of mαCD-mediated lipid exchange methods to study bilayer coupling in asymmetric cell membranes, opening a new door for examining by ImFCS.

## Developing the toolbox to study transbilayer interactions in live cells

We described in this *Perspective* general methods to investigate how dynamic membrane organization participates in transbilayer coupling (Fig. 1). Crosslinking of IgE-FcεRI (Ag-IgE-FcεRI; ^18, 37^) and GM1 (Ab-CTxB-GM1; ^48^) to induce transbilayer interactions have been established. Cholesterol extraction also appears to perturb these interactions but is complicated by cholesterol flipping between leaflets and other possible cellular impacts. The recently introduced lipid exchange (LEX) method for manipulating outer leaflet lipids in live cells ^50, 51^ offers exciting new opportunities for exploration. We also described herein a quantitative ImFCS-based platform using Lo- and Ld-preferring lipid probes to measure changes in the lipid-phase-like organization in the inner leaflet that accompany transbilayer interactions with exceptional sensitivity. If a particular treatment stabilizes Lo/Ld phase-like separation (e.g., Ag-IgE-FcεRI), the diffusion of an Lo-preferring probe is expected to decrease while that of an Ld-preferring probe increases (Fig. 3A). If a treatment causes global changes, impacting both Lo- and Ld-like regions similarly, the trend of diffusion change may be in the same direction for both Lo- and Ld-preferring probes (Figs. 3B,C). We emphasize that employing ImFCS to determine quantitative changes in membrane lipid organization is not limited to monitoring transbilayer coupling; these diffusion measurements are sensitive to effects of treatments that affect one or both leaflets (e.g., cholesterol depletion/repletion ^76^ and inhibition of actin polymerization ^7, 46^).

We have primarily exploited the exceptional data statistics of ImFCS obtained from multiple cells to detect subtle changes in the inner leaflet phase-like separation due to transbilayer interactions (Figs 2, 3). The raw image stacks recorded for ImFCS analysis are spatially resolved, and this spatial information can be extracted from the data set. For example, the diffusion maps derived from ImFCS measurements (e.g., Fig. 2A) was previously used to directly monitor the formation and time-dependent growth of micron-scale, solid-like peptide-lipid aggregates after addition of an amyloid peptide to live cells ^105^. More challenging is discerning phase-like separation in resting plasma membranes, which is nanoscopic such that individual Lo-like nanodomains are much smaller (<20 nm) than the spatial resolution of ImFCS-derived diffusion maps (Px unit = 320 nm in our measurements). However, emerging hypotheses, based on experiment and theory, posit that nanodomains may be connected by other membrane associated components (e.g., cortical actin cytoskeleton ^106, 107^) creating ordered membrane regions at length scales much longer than the size of individual nanodomains. Such long-range ordered structures would manifest non-homogenous diffusion maps at diffraction-limited spatial resolution. For example, actin-dense membrane regions may be expected to be more concentrated with nanodomains than are the actin-poor regions.

Two independent studies recently demonstrated these possibilities. Our ImFCS data from RBL plasma membranes showed that the diffusion maps of several lipid probes are not normally distributed ^7, 37^. However, distribution of diffusion coefficients of these probes may be simply fitted with two-component Gaussian model, indicating that there are two populations of Px units characterized by slower and faster diffusion, respectively. Px units with slower diffusing lipid probes correspond to stronger probe interactions with nanodomains. This may be due to higher density of these nanodomains in these Px units. Conversely, lower density of nanodomains in Px units corresponds to faster diffusion. Mashanov and colleagues made similar observations from their SPT measurements on fluorescently labeled M2 muscarinic acetylcholine receptor in live cells ^108^. In this study, the ventral plasma membrane is segmented into multiple 1-2 μm quadrats, and all SPT trajectories of a given quadrat are averaged to obtain an average diffusion coefficient of that quadrat. Repeating this process for all quadrats yields a diffusion map of the ventral membrane with 1-2 μm resolution. Interestingly, the diffusion maps of M2 acetylcholine receptor are not normally distributed in some cell types. Notably, M2 receptor was previously shown to undergo free diffusion in live cell membranes and therefore non-normal diffusion distribution reflects the presence of interaction-rich (slow diffusion) and interaction-poor (fast diffusion) membrane regions at 1-2 μm length scale. If nanodomains (and other membrane heterogeneity features) are randomly distributed throughout the cell membrane a normal distribution of diffusion (at 320-2000 nm length scale) is expected. These two studies therefore indicate a new level of membrane heterogeneity at much longer length scale than what is generally considered for membrane domains (20-400 nm) ^109^. Further insights on length scales of heterogeneity may also be gained from spatial correlation analyses (e.g., spot-variation FCS (svFCS) ^7, 110, 111^, image mean squared displacement (iMSD) ^66^, and pair-correlation function (pCF) ^67, 68^ analyses) applied on the same raw image stacks from ImFCS measurements. In future studies, it will be interesting to evaluate how various strategies of manipulating transbilayer interactions impact the non-normal distribution of long-range diffusion.

The transbilayer coupling modes presented herein (‘outside-in’, Fig. 1) offer manipulation of membrane properties with varying levels of experimental control. We described only one treatment condition for each category, and the impact of each (and other manipulations) can be characterized at a higher level of definition. In the case of Ag-IgE-FcεRI, for example, bivalent and trivalent Ag (instead of heterogeneous multivalent Ag) can be used to investigate which features of crosslinking/clustering are critical for transmembrane signaling^112^. The ImFCS platform can also be used to evaluate ‘inside out’ transbilayer interactions. For example, the phase-like properties in the outer leaflet can be measured before and after changing inner leaflet composition, such as by incorporating PS lipids of different acyl chain properties ^15^. For this purpose, it will be necessary to monitor and compare diffusion of a suitable pair of outer leaflet lipid probes that are Lo- and Ld-preferring. Interpretating ImFCS measurements will be further enhanced by complementary biophysical techniques. These include fluorescence anisotropy ^113^ and lifetime imaging microscopy of lipid probes ^6^ or polarity sensitive probes ^4^ in the inner and outer leaflet under defined stimulating or perturbing conditions.

The LEX methodology, which enables a wide range of lipids to be tested can be further explored and refined. It will be interesting to examine sphingomyelins with variable chain length because sphingomyelins are among the most abundant outer leaflet lipids, and they interact more strongly with cholesterol than their phospholipid counterparts ^114^. In addition, methods to exchange gangliosides and ceramides should be developed. Complications, such as effects on cytoskeletal organization, should also be evaluated. Ultimately, it will be important to determine lipid composition (including cholesterol) in each leaflet separately after LEX with different types of lipids. This is necessary to define the link between transbilayer coupling and lipid composition imposed by LEX, which presumably does not change significantly for Ag-IgE-FcεRI and Ab-CTxB-GM1 mediated transbilayer interactions. Quantification of outer lipid composition may be accomplished with the digestion method and lipidomics analysis^4^, or by lipid exchange and analysis of lipids removed from the cells ^50^, as previously reported.

For interpretation of ImFCS and other physical measurements in the biological context, the functional consequences of modulating transbilayer interactions must be quantified. Ag-crosslinking of IgE-FcεRI triggers a measurable signaling cascade in mast cells ^115^ that initiates allergic and inflammatory responses physiologically. Likewise, artificial cross-linking of GM1, Ab-CTxB-GM1, is sufficient to activate mitogen-activated protein kinase (MAPK) pathway ^48^ which is central to cell differentiation and proliferation. It will be interesting to see, for example, if the LEX with saturated lipids induces MAPK activity. In this regard, we note that LEX with saturated and unsaturated lipids was recently shown to respectively increase and decrease net intracellular phosphorylation of ligand-bound insulin receptor ^116^. Beyond transmembrane interactions underlying membrane biophysical properties, we expect the LEX method to be useful for evaluating the participation of different types of lipids in other plasma membrane dependent processes such as host-pathogen interactions, transmembrane signaling, and membrane trafficking.

## ACKNOWLEDGEMENTS

This work is supported by National Institute of General Medical Sciences (NIGMS) Grant R01GM117552. The content is solely the responsibility of the authors and does not necessarily represent the official views of NIGMS or NIH.

## REFERENCES

1. Bretscher, M. S., Asymmetrical lipid bilayer structure for biological membranes. Nat New Biol 1972, 236 (61), 11–2.

2. Doktorova, M.; Symons, J. L.; Levental, I., Structural and functional consequences of reversible lipid asymmetry in living membranes. Nature chemical biology 2020, 16 (12), 1321–1330.

3. Kobayashi, T.; Menon, A. K., Transbilayer lipid asymmetry. Curr Biol 2018, 28, PR386–R391.

4. Lorent, J. H.; Levental, K. R.; Ganesan, L.; Rivera-Longsworth, G.; Sezgin, E.; Doktorova, M. D.; Lyman, E.; Levental, I., Plasma membranes are asymmetric in lipid unsaturation, packing and protein shape. Nature chemical biology 2020.

5. Leventis, R.; Silvius, J., Use of Cyclodextrins to Monitor Transbilayer Movement and Differential Lipid Affinities of Cholesterol. Biophysical journal 2001, 81, 2257–2267.

6. Gupta, A.; Korte, T.; Herrmann, A.; Wohland, T., Plasma membrane asymmetry of lipid organization: fluorescence lifetime microscopy and correlation spectroscopy analysis. Journal of lipid research 2020, 61 (2), 252–266.

7. Bag, N.; Holowka, D. A.; Baird, B. A., Imaging FCS delineates subtle heterogeneity in plasma membranes of resting mast cells. Molecular biology of the cell 2020, 31 (7), 709–723.

8. Sezgin, E.; Levental, I.; Mayor, S.; Eggeling, C., The mystery of membrane organization: composition, regulation and roles of lipid rafts. Nature reviews. Molecular cell biology 2017, 18 (6), 361–374.

9. Levental, I.; Levental, K. R.; Heberle, F. A., Lipid Rafts: Controversies Resolved, Mysteries Remain. Trends in cell biology 2020.

10. Levental, I.; Veatch, S., The Continuing Mystery of Lipid Rafts. J Mol Biol 2016, 428 (24 Pt A), 4749–4764.

11. Shaw, T.; Ghosh, S.; Veatch, S., Critical Phenomena in Plasma Membrane Organization and Function. Annu Rev Phys Chem 2020.

12. Brown, D. A.; Rose, J. K., Sorting of GPI-anchored proteins to glycolipid-enriched membrane subdomains during transport to the apical cell surface. Cell 1992, 68 (3), 533–44.

13. Eggeling, C.; Ringemann, C.; Medda, R.; Schwarzmann, G.; Sandhoff, K.; Polyakova, S.; Belov, V. N.; Hein, B.; von Middendorff, C.; Schonle, A.; Hell, S. W., Direct observation of the nanoscale dynamics of membrane lipids in a living cell. Nature 2009, 457 (7233), 1159–62.

14. Owen, D. M.; Williamson, D. J.; Magenau, A.; Gaus, K., Sub-resolution lipid domains exist in the plasma membrane and regulate protein diffusion and distribution. Nature communications 2012, 3, 1256.

15. Raghupathy, R.; Anilkumar, A. A.; Polley, A.; Singh, P. P.; Yadav, M.; Johnson, C.; Suryawanshi, S.; Saikam, V.; Sawant, S. D.; Panda, A.; Guo, Z.; Vishwakarma, R. A.; Rao, M.; Mayor, S., Transbilayer lipid interactions mediate nanoclustering of lipid-anchored proteins. Cell 2015, 161 (3), 581–94.

16. Nicovich, P. R.; Kwiatek, J. M.; Ma, Y.; Benda, A.; Gaus, K., FSCS Reveals the Complexity of Lipid Domain Dynamics in the Plasma Membrane of Live Cells. Biophysical journal 2018, 114 (12), 2855–2864.

17. Mueller, V.; Ringemann, C.; Honigmann, A.; Schwarzmann, G.; Medda, R.; Leutenegger, M.; Polyakova, S.; Belov, V. N.; Hell, S. W.; Eggeling, C., STED Nanoscopy Reveals Molecular Details of Cholesterol- and Cytoskeleton-Modulated Lipid Interactions in Living Cells. Biophysical journal 2011, 101 (7), 1651–1660.

18. Shelby, S. A.; Veatch, S. L.; Holowka, D. A.; Baird, B. A., Functional nanoscale coupling of Lyn kinase with IgE-FcepsilonRI is restricted by the actin cytoskeleton in early antigen-stimulated signaling. Molecular biology of the cell 2016, 27 (22), 3645–3658.

19. Stone, M. B.; Shelby, S. A.; Nunez, M. F.; Wisser, K.; Veatch, S. L., Protein sorting by lipid phase-like domains supports emergent signaling function in B lymphocyte plasma membranes. eLife 2017, 6.

20. Honigmann, A.; Mueller, V.; Ta, H.; Schoenle, A.; Sezgin, E.; Hell, S. W.; Eggeling, C., Scanning STED-FCS reveals spatiotemporal heterogeneity of lipid interaction in the plasma membrane of living cells. Nature communications 2014, 5, 5412.

21. Saka, S. K.; Honigmann, A.; Eggeling, C.; Hell, S. W.; Lang, T.; Rizzoli, S. O., Multi-protein assemblies underlie the mesoscale organization of the plasma membrane. Nature communications 2014, 5, 4509.

22. Koster, D. V.; Mayor, S., Cortical actin and the plasma membrane: inextricably intertwined. Current opinion in cell biology 2016, 38, 81–9.

23. Honigmann, A.; Pralle, A., Compartmentalization of the Cell Membrane. J Mol Biol 2016, 428 (24 Pt A), 4739–4748.

24. Brown, D. A.; London, E., Structure and origin of ordered lipid domains in biological membranes. The Journal of membrane biology 1998, 164 (2), 103–14.

25. Kusumi, A.; Fujiwara, T. K.; Tsunoyama, T. A.; Kasai, R. S.; Liu, A. A.; Hirosawa, K. M.; Kinoshita, M.; Matsumori, N.; Komura, N.; Ando, H.; Suzuki, K. G. N., Defining raft domains in the plasma membrane. Traffic 2019.

26. Li, G.; Wang, Q.; Kakuda, S.; London, E., Nanodomains can persist at physiologic temperature in plasma membrane vesicles and be modulated by altering cell lipids. Journal of lipid research 2020, 61 (5), 758–766.

27. Wang, T.-Y.; Silvius, J. R., Cholesterol Does Not Induce Segregation of Liquid-Ordered Domains in Bilayers Modeling the Inner Leaflet of the Plasma Membrane. Biophysical journal 2001, 81, 2762–2773.

28. Baumgart, T.; Hammond, A. T.; Sengupta, P.; Hess, S. T.; Holowka, D. A.; Baird, B. A.; Webb, W. W., Large-scale fluid/fluid phase separation of proteins and lipids in giant plasma membrane vesicles. Proceedings of the National Academy of Sciences of the United States of America 2007, 104 (9), 3165–70.

29. St Clair, J. W.; Kakuda, S.; London, E., Induction of Ordered Lipid Raft Domain Formation by Loss of Lipid Asymmetry. Biophysical journal 2020, 119 (3), 483–492.

30. Fujimoto, T.; Parmryd, I., Interleaflet Coupling, Pinning, and Leaflet Asymmetry-Major Players in Plasma Membrane Nanodomain Formation. Frontiers in cell and developmental biology 2016, 4, 155.

31. Corradi, V.; Sejdiu, B. I.; Mesa-Galloso, H.; Abdizadeh, H.; Noskov, S. Y.; Marrink, S. J.; Tieleman, D. P., Emerging Diversity in Lipid-Protein Interactions. Chemical reviews 2019, 119 (9), 5775–5848.

32. Holowka, D.; Baird, B., Roles for lipid heterogeneity in immunoreceptor signaling. Biochimica et biophysica acta 2016, 1861 (8 Pt B), 830–836.

33. Mitra, E. D.; Whitehead, S. C.; Holowka, D.; Baird, B.; Sethna, J. P., Computation of a Theoretical Membrane Phase Diagram and the Role of Phase in Lipid-Raft-Mediated Protein Organization. J Phys Chem B 2018, 122 (13), 3500–3513.

34. Barua, D.; Goldstein, B., A Mechanistic Model of Early FceRI Signaling: Lipid Rafts and the Question of Protection from Dephosphorylation. PloS one 2012, 7 (12), e51669.

35. Field, K. A.; Holowka, D.; Baird, B., FceRI-mediated recruitment of p53/56^lyn^ to detergent-resistant membrane domains accompanies cellular signaling. Proceedings of the National Academy of Sciences of the United States of America 1995, 92, 9201–9205.

36. Davey, A. M.; Krise, K. M.; Sheets, E. D.; Heikal, A. A., Molecular perspective of antigen-mediated mast cell signaling. J Biol Chem 2008, 283 (11), 7117–27.

37. Bag, N.; Wagenknecht-Wiesner, A.; Lee, A.; Shi, S. M.; Holowka, D. A.; Baird, B. A., Lipid-based and protein-based interactions synergize transmembrane signaling stimulated by antigen clustering of IgE receptors. Proceedings of the National Academy of Sciences of the United States of America 2021, 118 (35).

38. Urbančič, I.; Schiffelers, L.; Jenkins, E.; Gong, W.; Santos, A. M.; Schneider, F.; O’Brien-Ball, C.; Vuong, M. T.; Ashman, N.; Sezgin, E.; Eggeling, C., Aggregation and immobilisation of membrane proteins interplay with local lipid order in the plasma membrane of T cells. bioRxiv 2020.

39. Sohn, H. W.; Tolar, P.; Jin, T.; Pierce, S. K., Fluorescence resonance energy transfer in living cells reveals dynamic membrane changes in the initiation of B cell signaling. Proceedings of the National Academy of Sciences of the United States of America 2006, 103 (21), 8143–8.

40. Gaus, K.; Chklovskaia, E.; Fazekas de St Groth, B.; Jessup, W.; Harder, T., Condensation of the plasma membrane at the site of T lymphocyte activation. The Journal of cell biology 2005, 171 (1), 121–31.

41. Kiessling, V.; Wan, C.; Tamm, L. K., Domain coupling in asymmetric lipid bilayers. Biochimica et biophysica acta 2009, 1788 (1), 64–71.

42. Sarmento, M. J.; Hof, M.; Sachl, R., Interleaflet Coupling of Lipid Nanodomains - Insights From in vitro Systems. Frontiers in cell and developmental biology 2020, 8, 284.

43. London, E., Membrane Structure-Function Insights from Asymmetric Lipid Vesicles. Acc Chem Res 2019, 52 (8), 2382–2391.

44. Chiantia, S.; London, E., Acyl chain length and saturation modulate interleaflet coupling in asymmetric bilayers: effects on dynamics and structural order. Biophysical journal 2012, 103 (11), 2311–9.

45. Lin, Q.; London, E., Ordered raft domains induced by outer leaflet sphingomyelin in cholesterol-rich asymmetric vesicles. Biophysical journal 2015, 108 (9), 2212–22.

46. Gupta, A.; Muralidharan, S.; Torta, F.; Wenk, M. R.; Wohland, T., Long acyl chain ceramides govern cholesterol and cytoskeleton dependence of membrane outer leaflet dynamics. Biochim Biophys Acta Biomembr 2020, 1862 (3), 183153.

47. Arumugam, S.; Schmieder, S.; Pezeshkian, W.; Becken, U.; Wunder, C.; Chinnapen, D.; Ipsen, J. H.; Kenworthy, A. K.; Lencer, W.; Mayor, S.; Johannes, L., Ceramide structure dictates glycosphingolipid nanodomain assembly and function. Nature communications 2021, 12 (1).

48. Koyama-Honda, I.; Fujiwara, T. K.; Kasai, R. S.; Suzuki, K. G. N.; Kajikawa, E.; Tsuboi, H.; Tsunoyama, T. A.; Kusumi, A., High-speed single-molecule imaging reveals signal transduction by induced transbilayer raft phases. Journal of Cell Biology 2020, 219 (12).

49. Dinic, J.; Ashrafzadeh, P.; Parmryd, I., Actin filaments attachment at the plasma membrane in live cells cause the formation of ordered lipid domains. Biochimica et biophysica acta 2013, 1828 (3), 1102–11.

50. Li, G.; Kim, J.; Huang, Z.; St Clair, J. R.; Brown, D. A.; London, E., Efficient replacement of plasma membrane outer leaflet phospholipids and sphingolipids in cells with exogenous lipids. Proceedings of the National Academy of Sciences of the United States of America 2016, 113 (49), 14025–14030.

51. Li, G.; Kakuda, S.; Suresh, P.; Canals, D.; Salamone, S.; London, E., Replacing plasma membrane outer leaflet lipids with exogenous lipid without damaging membrane integrity. PLoS One 2019, 14 (10), e0223572.

52. Suresh, P.; London, E., Using cyclodextrin-induced lipid substitution to study membrane lipid and ordered membrane domain (raft) function in cells. Biochim Biophys Acta Biomembr 2021, 1864 (1), 183774.

53. Levental, I.; Byfield, F. J.; Chowdhury, P.; Gai, F.; Baumgart, T.; Janmey, P. A., Cholesterol-dependent phase separation in cell-derived giant plasma-membrane vesicles. The Biochemical journal 2009, 424 (2), 163–7.

54. Baumgart, T.; Hunt, G.; Farkas, E. R.; Webb, W. W.; Feigenson, G. W., Fluorescence probe partitioning between Lo/Ld phases in lipid membranes. Biochimica et biophysica acta 2007, 1768 (9), 2182–2194.

55. Heberle, F. A.; Doktorova, M.; Scott, H. L.; Skinkle, A. D.; Waxham, M. N.; Levental, I., Direct label-free imaging of nanodomains in biomimetic and biological membranes by cryogenic electron microscopy. Proc Natl Acad Sci 2020, 117 (33), 19943–19952.

56. Cornell, C. E.; Mileant, A.; Thakkar, N.; Lee, K. K.; Keller, S. L., Direct imaging of liquid domains in membranes by cryo-electron tomography. Proceedings of the National Academy of Sciences of the United States of America 2020, 117 (33), 19713–19719.

57. Varma, R.; Mayor, S., GPI-anchored proteins are organized in submicron domains at the cell surface. Nature 1998, 394 (6695), 798–801.

58. Sengupta, P.; Holowka, D.; Baird, B., Fluorescence resonance energy transfer between lipid probes detects nanoscopic heterogeneity in the plasma membrane of live cells. Biophysical journal 2007, 92 (10), 3564–74.

59. Shelby, S. A.; Castello-Serrano, I.; Wisser, K. C.; Levental, I.; Veatch, S. L., Membrane phase separation drives organization at B cell receptor clusters. BioRxiv 2021.

60. Sezgin, E.; Schwille, P., Fluorescence techniques to study lipid dynamics. Cold Spring Harbor perspectives in biology 2011, 3 (11), a009803.

61. Kenworthy, A. K.; Nichols, B. J.; Remmert, C. L.; Hendrix, G. M.; Kumar, M.; Zimmerberg, J.; Lippincott-Schwartz, J., Dynamics of putative raft-associated proteins at the cell surface. The Journal of cell biology 2004, 165 (5), 735–46.

62. Bacia, K.; Scherfeld, D.; Kahya, N.; Schwille, P., Fluorescence correlation spectroscopy relates rafts in model and native membranes. Biophysical journal 2004, 87 (2), 1034–43.

63. Sanchez, S. A.; Tricerri, M. A.; Gratton, E., Laurdan generalized polarization fluctuations measures membrane packing micro-heterogeneity in vivo. Proceedings of the National Academy of Sciences of the United States of America 2012, 109 (19), 7314–9.

64. Wawrezinieck, L.; Rigneault, H.; Marguet, D.; Lenne, P., Fluorescence correlation spectroscopy diffusion laws to probe the submicron cell membrane organization. Biophysical journal 2005, 89 (6), 4029–4042.

65. Yechiel, E.; Edidin, M., Micrometer-scale domains in fibroblast plasma membranes. The Journal of cell biology 1987, 105 (2), 755–60.

66. Di Rienzo, C.; Gratton, E.; Beltram, F.; Cardarelli, F., Fast spatiotemporal correlation spectroscopy to determine protein lateral diffusion laws in live cell membranes. Proceedings of the National Academy of Sciences of the United States of America 2013, 110 (30), 12307–12.

67. Digman, M. A.; Gratton, E., Imaging barriers to diffusion by pair correlation functions. Biophysical journal 2009, 97 (2), 665–73.

68. Sankaran, J.; Manna, M.; Guo, L.; Kraut, R.; Wohland, T., Diffusion, transport, and cell membrane organization investigated by imaging fluorescence cross-correlation spectroscopy. Biophysical journal 2009, 97 (9), 2630–9.

69. Manzo, C.; van Zanten, T. S.; Garcia-Parajo, M. F., Nanoscale fluorescence correlation spectroscopy on intact living cell membranes with NSOM probes. Biophysical journal 2011, 100 (2), L8–10.

70. Winkler, P. M.; Regmi, R.; Flauraud, V.; Brugger, J.; Rigneault, H.; Wenger, J.; Garcia-Parajo, M. F., Optical Antenna-Based Fluorescence Correlation Spectroscopy to Probe the Nanoscale Dynamics of Biological Membranes. The journal of physical chemistry letters 2018, 9 (1), 110–119.

71. Fujiwara, T.; Ritchie, K.; Murakoshi, H.; Jacobson, K.; Kusumi, A., Phospholipids undergo hop diffusion in compartmentalized cell membrane. The Journal of cell biology 2002, 157 (6), 1071–81.

72. Sahl, S. J.; Leutenegger, M.; Hilbert, M.; Hell, S. W.; Eggeling, C., Fast molecular tracking maps nanoscale dynamics of plasma membrane lipids. Proceedings of the National Academy of Sciences of the United States of America 2010, 107 (15), 6829–34.

73. Krieger, J. W.; Singh, A. P.; Bag, N.; Garbe, C. S.; Saunders, T. E.; Langowski, J.; Wohland, T., Imaging fluorescence (cross-) correlation spectroscopy in live cells and organisms. Nature protocols 2015, 10 (12), 1948–74.

74. Wohland, T. https://www.dbs.nus.edu.sg/lab/BFL/APEER_ImagingFCS.html and https://www.dbs.nus.edu.sg/lab/BFL/imfcs_image_j_plugin.html.

75. Bag, N.; Wohland, T., Imaging fluorescence fluctuation spectroscopy: new tools for quantitative bioimaging. Annu Rev Phys Chem 2014, 65, 225–48.

76. Bag, N.; Huang, S.; Wohland, T., Plasma Membrane Organization of Epidermal Growth Factor Receptor in Resting and Ligand-Bound States. Biophysical journal 2015, 109 (9), 1925–36.

77. Ng, X. W.; Teh, C.; Korzh, V.; Wohland, T., The secreted signaling protein Wnt3 is associated with plasma membrane lipid domains *in vivo:* a SPIM-FCS study. Biophysical journal 2016, 111, 418–429.

78. Bag, N.; Huang, S.; Wohland, T., Investigating the Dynamics and Organization of Membrane Proteins and Lipids by Imaging Fluorescence Correlation Spectroscopy. In Membrane Organization and Dynamics, Springer Series in Biophysics: 2017; Vol. 20, pp 113–145.

79. Edwald, E.; Stone, M. B.; Gray, E. M.; Wu, J.; Veatch, S. L., Oxygen depletion speeds and simplifies diffusion in HeLa cells. Biophysical journal 2014, 107 (8), 1873–84.

80. Menon, A. K.; Holowka, D.; Webb, W. W.; Baird, B., Cross-Linking of Receptor-Bound Ige to Aggregates Larger Than Dimers Leads to Rapid Immobilization. Journal of Cell Biology 1986, 102 (2), 541–550.

81. Harder, T.; Scheiffele, P.; Verkade, P.; Simons, K., Lipid Domain Structure of the Plasma Membrane revaled by patching of membrane components. J Cell Bio 1998, 141 (14), 929–942.

82. van Zanten, T. S.; Gomez, J.; Manzo, C.; Cambi, A.; Buceta, J.; Reigada, R.; Garcia-Parajo, M. F., Direct mapping of nanoscale compositional connectivity on intact cell membranes. Proceedings of the National Academy of Sciences of the United States of America 2010, 107 (35), 15437–42.

83. Andrews, N. L.; Lidke, K. A.; Pfeiffer, J. R.; Burns, A. R.; Wilson, B. S.; Oliver, J. M.; Lidke, D. S., Actin restricts FcepsilonRI diffusion and facilitates antigen-induced receptor immobilization. Nat Cell Biol 2008, 10 (8), 955–63.

84. Kabbani, A. M.; Raghunathan, K.; Lencer, W. I.; Kenworthy, A. K.; Kelly, C. V., Structured clustering of the glycosphingolipid GM1 is required for membrane curvature induced by cholera toxin. Proceedings of the National Academy of Sciences of the United States of America 2020, 117 (26), 14978–14986.

85. Brown, D. A.; London, E., Functions of lipid rafts in biological membranes. Annual review of cell and developmental biology 1998, 14, 111–36.

86. Pike, L. J., Rafts defined: a report on the Keystone Symposium on Lipid Rafts and Cell Function. Journal of lipid research 2006, 47 (7), 1597–8.

87. Zidovetzki, R.; Levitan, I., Use of cyclodextrins to manipulate plasma membrane cholesterol content: evidence, misconceptions and control strategies. Biochimica et biophysica acta 2007, 1768 (6), 1311–24.

88. Amundson, D. M.; Zhou, M., Fluorometric method for the enzymatic determination of cholesterol. Journal of Biochemical and Biophysical Methods 1999, 38, 43–52.

89. Kim, J. H.; Singh, A.; Del Poeta, M.; Brown, D. A.; London, E., The effect of sterol structure upon clathrin-mediated and clathrin-independent endocytosis. Journal of cell science 2017, 130 (16), 2682–2695.

90. Bag, N.; Yap, D. H. X.; Wohland, T., Temperature dependence of diffusion in model and live cell membranes characterized by imaging fluorescence correlation spectroscopy. Biochimica et Biophysica Acta (BBA) - Biomembranes 2014, 1838, 802–813.

91. Nishimura, S. Y.; Vrljic, M.; Klein, L. O.; McConnell, H. M.; Moerner, W. E., Cholesterol depletion induces solid-like regions in the plasma membrane. Biophysical journal 2006, 90 (3), 927–38.

92. Mahammad, S.; Dinic, J.; Adler, J.; Parmryd, I., Limited cholesterol depletion causes aggregation of plasma membrane lipid rafts inducing T cell activation. Biochimica et biophysica acta 2010, 1801 (6), 625–34.

93. Steck, T. L.; Lange, Y., Transverse distribution of plasma membrane bilayer cholesterol: Picking sides. Traffic 2018, 19 (10), 750–760.

94. Liu, S. L.; Sheng, R.; Jung, J. H.; Wang, L.; Stec, E.; O’Connor, M. J.; Song, S.; Bikkavilli, R. K.; Winn, R. A.; Lee, D.; Baek, K.; Ueda, K.; Levitan, I.; Kim, K. P.; Cho, W., Orthogonal lipid sensors identify transbilayer asymmetry of plasma membrane cholesterol. Nature chemical biology 2017, 13 (3), 268–274.

95. Das, A.; Brown, M. S.; Anderson, D. D.; Goldstein, J. L.; Radhakrishnan, A., Three pools of plasma membrane cholesterol and their relation to cholesterol homeostasis. eLife 2014, 3.

96. Buwaneka, P.; Ralko, A.; Liu, S. L.; Cho, W., Evaluation of the available cholesterol concentration in the inner leaflet of the plasma membrane of mammalian cells. Journal of lipid research 2021, 62, 100084.

97. Courtney, K. C.; Fung, K. Y.; Maxfield, F. R.; Fairn, G. D.; Zha, X., Comment on ‘Orthogonal lipid sensors identify transbilayer asymmetry of plasma membrane cholesterol’. eLife 2018, 7.

98. Mondal, M.; Mesmin, B.; Mukherjee, S.; Maxfield, F. R., Sterols are mainly in the cytoplasmic leaflet of the plasma membrane and the endocytic recycling compartment in CHO cells. Molecular biology of the cell 2009, 20 (2), 581–8.

99. van Meer, G.; Voelker, D. R.; Feigenson, G. W., Membrane lipids: where they are and how they behave. Nature Reviews Molecular Cell Biology 2008, 9 (2), 112–124.

100. Sheets, E. D.; Holowka, D.; Baird, B., Critical Role for Cholesterol in Lyn-mediated Tyrosine Phosphorylation of FceRI and Their Association with Detergent-resistant Membranes. The Journal of cell biology 1999, 145 (877-887).

101. Kornberg, R. D.; McConnell, H. M., Inside-outside transitions of phospholipids in vesicle membranes. Biochemistry-Us 1971, 10 (7), 1111–20.

102. Huang, Z.; London, E., Effect of cyclodextrin and membrane lipid structure upon cyclodextrin-lipid interaction. Langmuir: the ACS journal of surfaces and colloids 2013, 29 (47), 14631–8.

103. Wang, Q.; London, E., Lipid Structure and Composition Control Consequences of Interleaflet Coupling in Asymmetric Vesicles. Biophysical journal 2018, 115 (4), 664–678.

104. Kakuda, S.; Suresh, P.; Li, G.; London, E., Loss of plasma membrane lipid asymmetry can induce ordered domain (raft) formation. Journal of lipid research 2021, 63 (1), 100155.

105. Bag, N.; Ali, A.; Chauhan, V. S.; Wohland, T.; Mishra, A., Membrane destabilization by monomeric hIAPP observed by imaging fluorescence correlation spectroscopy. Chemical communications 2013, 49 (80), 9155–9157.

106. Machta, B. B.; Papanikolaou, S.; Sethna, J. P.; Veatch, S. L., Minimal model of plasma membrane heterogeneity requires coupling cortical actin to criticality. Biophysical journal 2011, 100 (7), 1668–77.

107. Honigmann, A.; Sadeghi, S.; Keller, J.; Hell, S. W.; Eggeling, C.; Vink, R., A lipid bound actin meshwork organizes liquid phase separation in model membranes. eLife 2014, 3, e01671.

108. Mashanov, G. I.; Nenasheva, T. A.; Mashanova, A.; Lape, R.; Birdsall, N. J. M.; Sivilotti, L.; Molloy, J. E., Heterogeneity of cell membrane structure studied by single molecule tracking. Faraday Discuss 2021.

109. Kusumi, A.; Fujiwara, T. K.; Chadda, R.; Xie, M.; Tsunoyama, T. A.; Kalay, Z.; Kasai, R. S.; Suzuki, K. G., Dynamic organizing principles of the plasma membrane that regulate signal transduction: commemorating the fortieth anniversary of Singer and Nicolson’s fluid-mosaic model. Annual review of cell and developmental biology 2012, 28, 215–250.

110. Bag, N.; Sankaran, J.; Paul, A.; Kraut, R. S.; Wohland, T., Calibration and limits of camera-based fluorescence correlation spectroscopy: a supported lipid bilayer study. Chemphyschem: a European journal of chemical physics and physical chemistry 2012, 13 (11), 2784–94.

111. Veerapathiran, S.; Wohland, T., The imaging FCS diffusion law in the presence of multiple diffusive modes. Methods 2017.

112. Holowka, D.; Sil, D.; Torigoe, C.; Baird, B., Insights into immunoglobulin E receptor signaling from structurally defined ligands. Immnological Reviews 217, 269–279.

113. Rao, M.; Mayor, S., Use of Forster’s resonance energy transfer microscopy to study lipid rafts. Biochimica et biophysica acta 2005, 1746 (3), 221–33.

114. Silvius, J. R., Role of cholesterol in lipid raft formation-lessons from lipid model systems. Biochimica et Biophysica Acta (BBA) - Biomembranes 2003, 1610, 174–183.

115. Rivera, J.; Gilfillan, A. M., Molecular regulation of mast cell activation. The Journal of allergy and clinical immunology 2006, 117 (6), 1214–25; quiz 1226.

116. Suresh, P.; Miller, W. T.; London, E., Phospholipid exchange shows insulin receptor activity is supported by both the propensity to form wide bilayers and ordered raft domains. J Biol Chem 2021, 297 (3), 101010.

117. Pralle, A.; Keller, P.; Florin, E.-L.; Hörbor, J. K. H., Sphingolipid-Cholesterol Rafts Diffuse as Small Entities in the Plasma Membrane of Mammalian Cells. The Journal of cell biology 2000, 148 (5), 997-1007.

118. Sengupta, P.; Hammond, A.; Holowka, D.; Baird, B., Structural determinants for partitioning of lipids and proteins between coexisting fluid phases in giant plasma membrane vesicles. Biochimica et biophysica acta 2008, 1778 (1), 20–32.

119. Pyenta, P. S.; Holowka, D.; Baird, B., Cross-correlation analysis of inner-leaflet-anchored green fluorescent protein co-redistributed with IgE receptors and outer leaflet lipid raft components. Biophysical journal 2001, 80 (5), 2120–32.

120. Rodgers, W., Making Membranes Green: Construction and Characterization of GFPFusion Proteins Targeted to Discrete Plasma Membrane Domains. BioTechniques 2002, 32, 1044–1051.

